# Whistle variability and social acoustic interactions in bottlenose dolphins

**DOI:** 10.1101/2024.10.15.618471

**Authors:** Faadil Mustun, Chiara Semenzin, Dean Rance, Emiliano Marachlian, Zohria-Lys Guillerm, Agathe Mancini, Inès Bouaziz, Elisabeth Fleck, Nadav Shashar, Gonzalo G. de Polavieja, Germán Sumbre

## Abstract

Bottlenose dolphins exhibit a sophisticated social structure, known as a fission-fusion society. To sustain this complex system, dolphins rely on a rich vocal repertoire: clicks exclusively used for echolocation, burst-pulse sounds associated with emotions during social interactions, and whistles, including signature whistles that serve as individual-specific identifiers (‘names’).

How dolphins maintain their complex social structure based only on a limited repertoire of sounds remains elusive. Previous studies suggest that contextual information can be transferred by the modulation of the whistles.

Here, we investigated the whistle variability using a comprehensive computational approach, and studied the structure of the interactions between the whistle variants. Using a unique large dataset, acquired in a natural environment, we observed that signature whistles exhibit variations in their frequency contours. Unsupervised clustering revealed that signature whistles could be classified into sub-categories (signature whistle variants). The existence of these categories, and their independence on the emitter dolphin, indicate that these variations are not random. Analysis of pairwise interactions between sub-categories revealed a clustered structure similar to that of their social hierarchy. Network analysis of this structure showed that whistle sub-categories had different functional roles: some acted as hubs, others as bridges, and certain were used for turn-taking between the main whistle categories. We also found that the dolphins emit signature whistles of their deceased mothers, a phenomenon only observed in human language.

Overall, these findings suggest that dolphins modulate signature whistles to transmit additional information beyond individual identity, and that they engage in "dialogue-like" acoustic interactions.

## Introduction

Bottlenose dolphins display a highly developed social structure that is crucial for their survival and well-being. This social structure is described as a fission-fusion society, in which the composition and splitting of groups dynamically changes over time according to the associated costs and benefits (1–8). For example, males form strong and long-lasting bonds with one or two other males, and second-order affiliations with other groups of males. Females usually regroup in larger pods assuring protection for their calves. These female and male pods can form superpods containing several hundred individuals. Dolphins can separate from their reference pods and spend several minutes to years with other pods or alone.

In order to maintain this complex social system, dolphins need to transmit elaborate information over long distances. In the sea, sound is the most efficient communication modality, capable of propagating up to ∼10 km (9, 10). The vocal repertoire of dolphins spans across a variety of pulse and pure tones that can be classified into three main categories (11, 12): clicks, burst pulse sounds, and whistles. Clicks are short duration broadband sounds emitted at low repetition rates. These signals are used for echolocation during navigation and foraging. Burst pulse sounds (BPS), are packets of closely spaced broadband clicks, produced at a higher rate than echolocation clicks. BPS are associated with courtship, dominance (13), and aggressive behaviors (14), suggesting that BPS play a role in communicating emotions during social interactions. Whistles are continuous narrow-band and frequency-modulated sounds that may contain multiple harmonics. Whistles are divided into two types: signature whistles (SWs) and non-signature whistles (NSWs). NSWs are thought to be employed in social contexts, but their role remains elusive (15). In contrast, SWs are frequency contours broadcasting the identity of the owner (12, 16). SWs develop during the first year of life, and then remain stable throughout the animal’s lifetime (17). Nevertheless, dolphins can modify the frequency range of their SWs to prevent interference in noisy environments (18, 19). Although each SW is predominantly vocalized by the owner, dolphins may also emit the SW of their conspecifics (vocal matching) (20–24). SWs represent a significant portion of the vocalization repertoire (38–70%) in the wild (25), and about 90% in isolation (25–27). Therefore, it was suggested that SWs act as “names” that can be used to transmit a dolphin’s identity or call conspecifics out of sight (vocal labeling). Vocal labeling has also been observed in elephants (28) and marmoset monkeys (29).

In dolphins, SWs may show certain variability. For example, during vocal mimicry (emitting the SW of a different dolphin) of SWs emitted by other individuals (23) or synthetic playbacked whistles (30), dolphins can introduce fine-scale frequency or duration modifications to the SW, while conserving the main frequency contour.

In a one-individual study, a captive bottlenose dolphin modulated the onset frequency of its SW depending on the experimental context (isolation vs. following correct choice in a discrimination task) (31). Moreover, analysis of the overall whistle population of different dolphin pods (Sardinia and Croatia), revealed different whistle repertoires that were associated with both social and behavioral contexts (e.g. feeding, traveling or group size) (32). More recently, it was observed that female bottlenose dolphins variate the maximum frequency and frequency range of their SWs depending on the presence or the absence of their calves (33). These studies suggest the hypothesis that identity and context may be encoded in the variability of the whistles.

Despite these findings, a comprehensive analysis of the whistle variability has never been done. Moreover, the structure of the interactions between the different whistle variants is unknown.

Here, we addressed these open questions using a computational approach to analyze a unique large dataset, acquired from a pod of dolphins, with a well-known social structure, living in a semi-natural environment (Dolphin Reef, Eilat) (34). We observed that the emitted SWs displayed consistent and specific variations of their frequency contour which could be classified in discrete groups (whistle sub-categories). In addition, these whistle variations could not be explained by the identity of the emitter. Thus, we suggest that the observed variability of the SWs carries additional information, beyond the identity of the owner. Then, we computed the structure of the whistle sub-categories, and used network analysis to uncover their role in acoustic communication. The pairwise interactions between the different sub-categories of whistles revealed a clustered structured network that resembles that of their social organization. Among these whistle sub-categories, some displayed different functional roles in the network. We thus argue that these whistle sub-categories are strong candidates for distinct communication units and may serve different communicative functions.

## Results

In order to obtain valuable insights into dolphin acoustic communication, it is necessary to follow and record the acoustic interactions of the same individuals over long periods of time. In our study, we recorded 224 hours of dolphin acoustic interactions during 115 days from the same 5 dolphins in the Dolphin reef, Eilat (Supplementary Fig.5a). From these recordings, we extracted 8566 whistles using a custom-made algorithm based on the variance of the spectrograms and their distance with respect to known whistle templates, based on dynamic time warping (DTW, Supplementary Fig.1, see Methods). The extracted whistles were then classified using ARTwarp (35). This clustering approach identified 10 main categories of whistles: 4 representing SWs of the dolphins born in the area of the Dolphin Reef (*Tursiops truncatus ponticus,* Fig.1a-c), one SW of a solitary dolphin that joined the pod of the Dolphin reef for a limited period of time (*Tursiops aduncus*, Fig.1b), and five NSWs (Fig.1c). Whistles that occurred only once in the dataset were not included in subsequent analyses. The 4 dolphins born in the reef are called Neo (the only male), Nikita, Nana and Luna (females). The solitary dolphin is called Yosefa (female, Supplementary Fig.5a).

### Whistle variability

We observed that, within each of the whistle main categories, the main frequency contours were conserved but they displayed variations in terms of frequency and duration (stretching and compression, Supplementary Fig. 2). To investigate whether this variability represents a random process or it rather displays a conserved structure, we re-cluster the whistle categories into sub-groups, using again the unsupervised algorithm ARTwarp (see Methods). The re-clustering showed that most of the whistles could, on average, be classified into 8.9±4.8 discrete groups (whistle sub-categories, Fig. 1).

**Figure 1.**
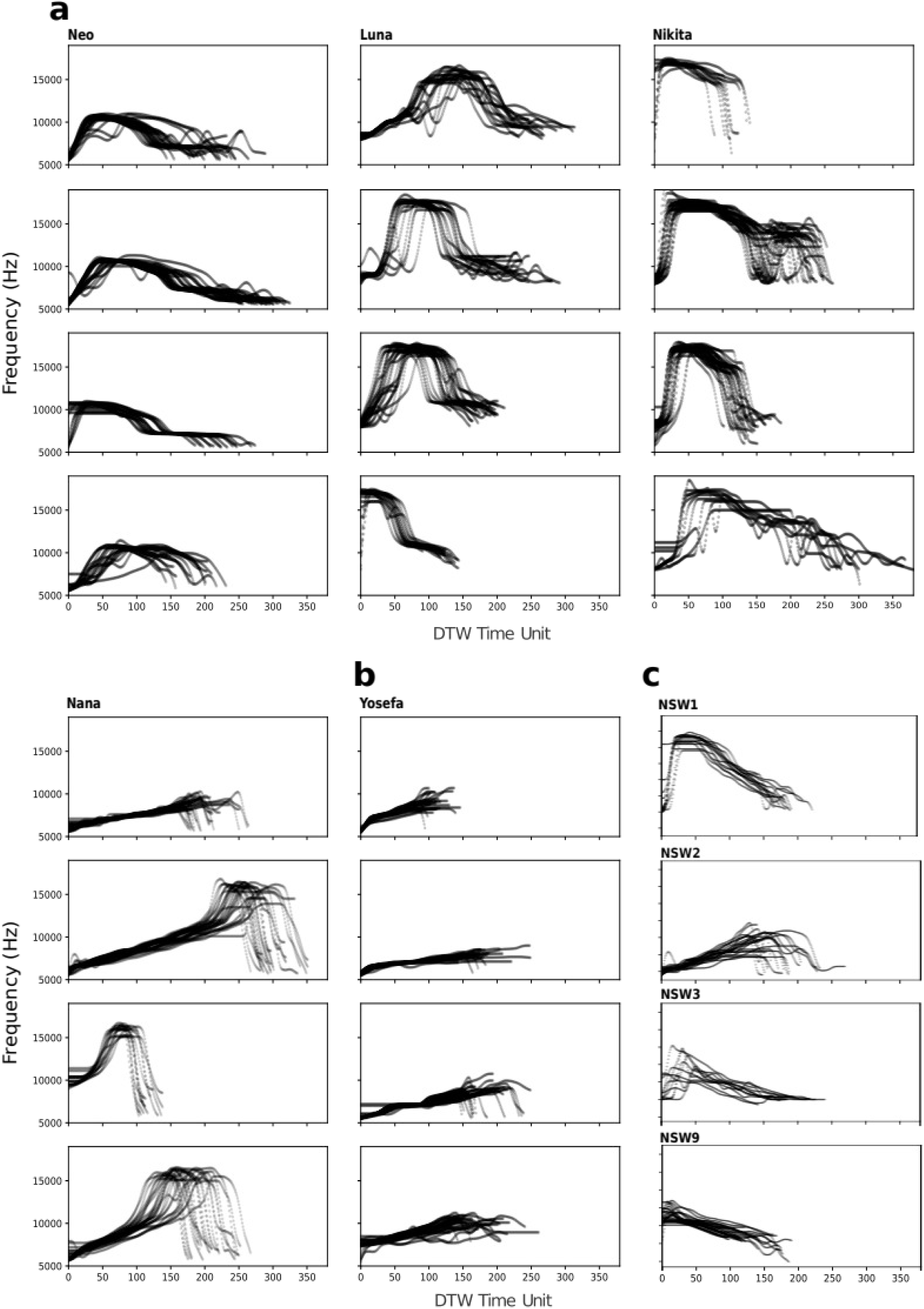
Whistles show variability of their frequency contours that can be classified in discrete sub-categories. a) Four example sub-categories of the SW_Neo, SW_Luna, SW_Nikita and SW_Nana. Each panel shows several superimposed frequency contours belonging to one sub-category. For visualization purposes, the whistles were aligned to a model whistle according to the DTW algorithm. b) As (a) for the dolphin that does not belong to the pod, Yosefa. c) As for (a) for SW_Dana (mother of Neo and Luna, deceased in 2013), SW_Shy (mother of Nikita and Nana, deceased in 2015), and NSW_3. Note the reduced variability within sub-categories, and the differences between sub-categories.

To study whether these frequency variations occur at specific areas of the frequency contour of the whistles, we used a distance measure based on DTW to align the whistles (a time-invariant shape alignment). Then, the whistles were re-clustered using K-Means to capture their different shapes (modulation patterns such as deletions and additions). While these clusters did not capture the full inventory of whistles, it showed that certain types of variability occurred at specific portions of the frequency contours. For example, Neo’s modulations were clearly concentrated at the top-most part of the SW and at its end (Fig.2a and Supplementary Fig.1d), while Nikita’s SW was mainly modulated in the region that directly precedes the end of the whistle (Fig. 2b). These results suggest that the variability of the whistles is not the result of a random process, as it can be classified into discrete groups, and it occurs at specific regions of the frequency contours.

**Figure 2.**
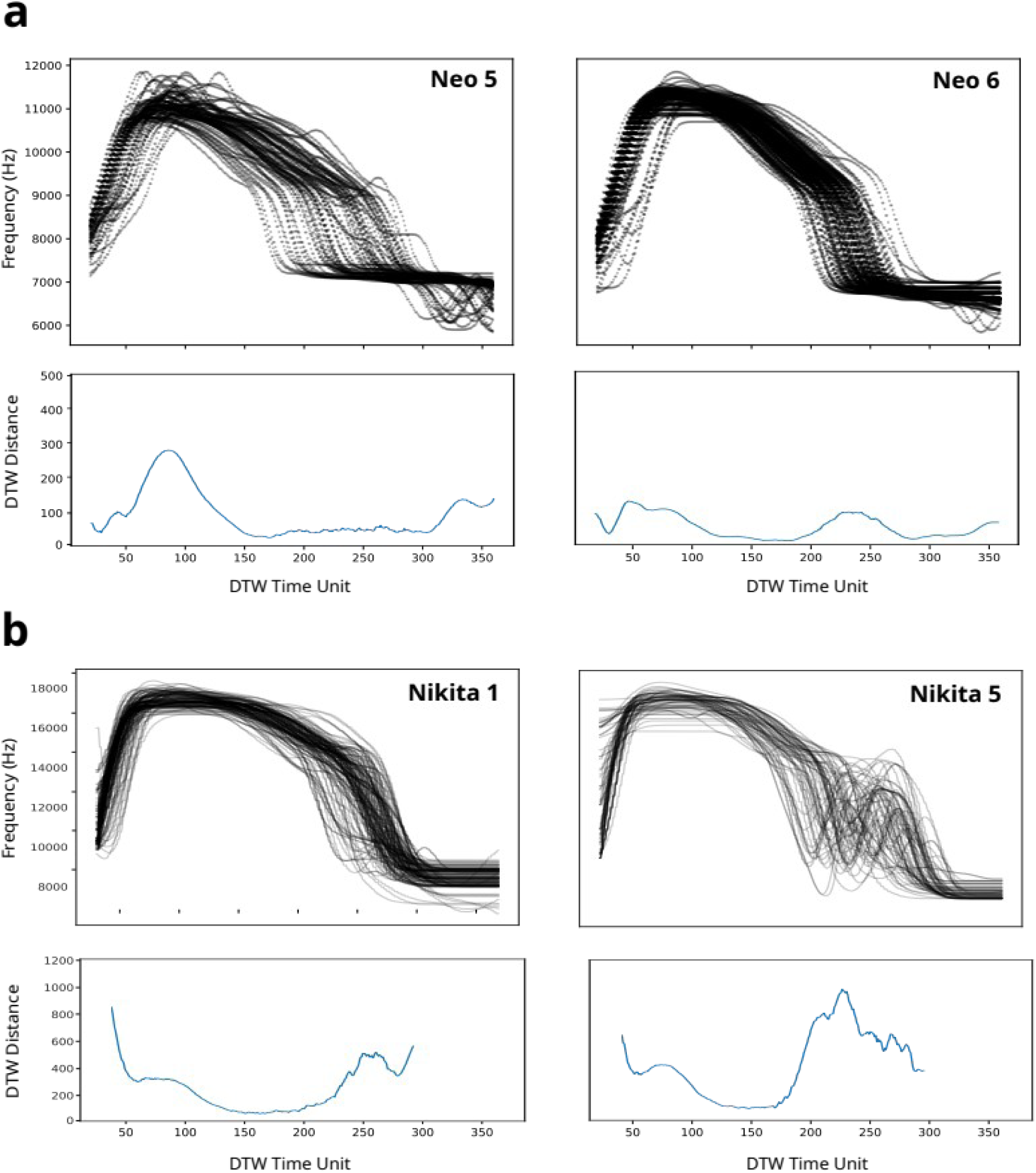
Certain whistle modulations occur at specific time points along the frequency contour of the whistles. a) Top: The superimposed frequency contours of two examples of SW_Neo sub-categories (Neo 5 and Neo 6). The frequency contours were aligned using DTW. Bottom: The mean distance between a whistle model and each of the frequency contours on top, at every DTW unit. The peaks represent the time points where the frequency contour of the whistles show the largest variability. b) As (a) for two SW_Nikita sub-categories. Note how certain frequency modulations occur at specific time points along the frequency contours.

Since dolphins can mimic the SWs of others, it is possible that the different SW sub-categories represent the dolphin that emitted the whistle (each dolphin emits the SWs with a distinctive modulation). To test this hypothesis, we analyzed the correspondence between the emitter identity and the variability in the whistles. For this purpose, we used a smaller dataset of 402 emitter-labeled SWs. This labeled dataset was obtained from simultaneous video and audio recordings, from which we extracted SWs when only one dolphin was in the surroundings (see Methods). We found that all SW main categories could be classified into more sub-categories than there are individuals in the pod. We also observed that a significant portion of the whistle sub-categories were emitted by more than one dolphin (from 26 whistle sub-categories, 17 involved more than one individual, n= 402, Fig.3). This suggests that the observed whistle variance can not be fully explained by the identity of the dolphin that emitted it.

**Figure 3.**
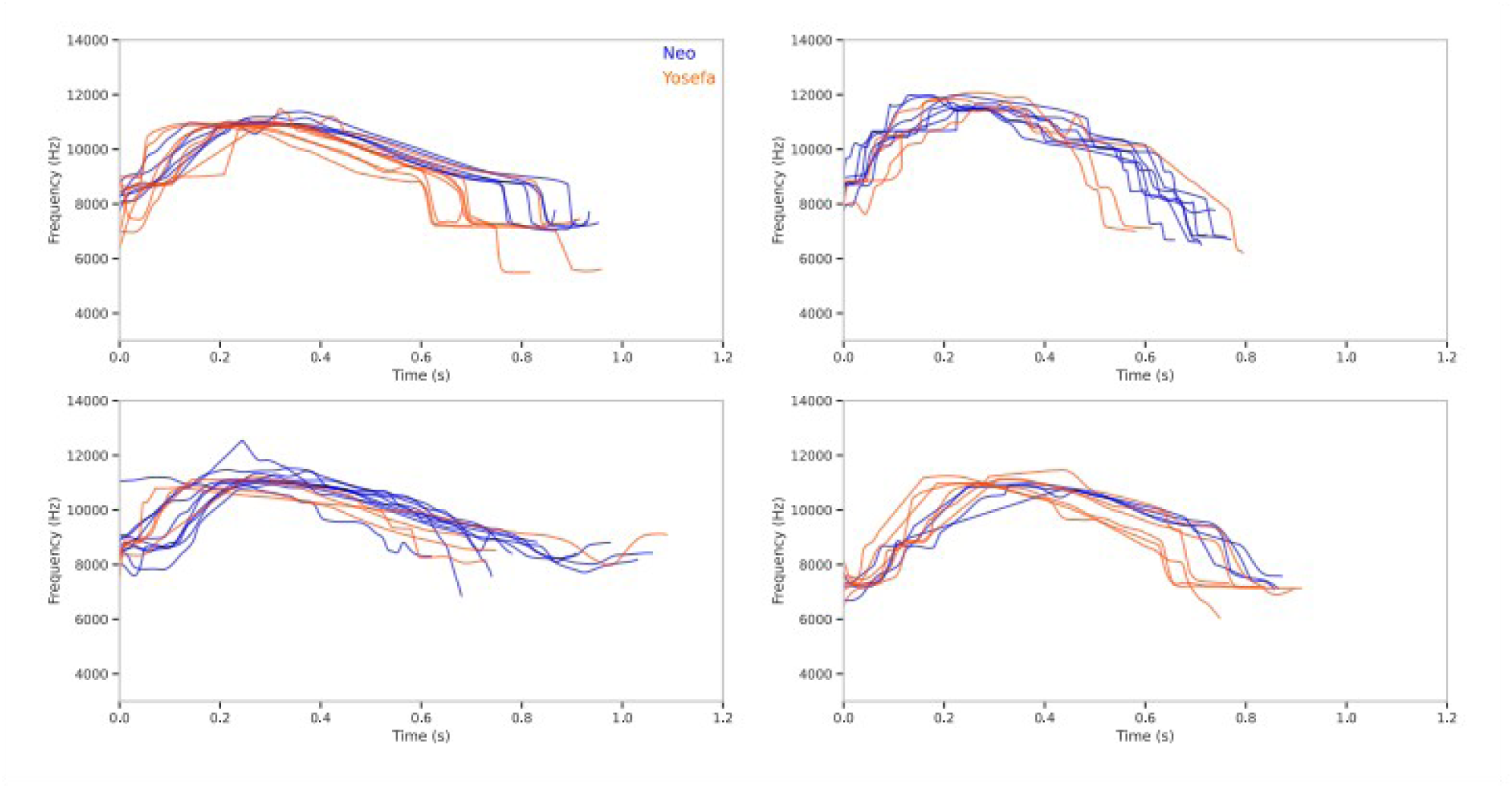
SW sub-categories do not represent different dolphin emitters. Examples of the frequency contours of 4 sub-categories of SW_Neo. Using the diarized dataset, we identified for each SW_Neo, the identity of the emitter. Blue: Neo (N=27). Red: Yosefa (N=21). Note that both emitters are present in each SW_Neo sub-category.

This hypothesis was also supported by the observation of connected sequences of whistles (series of whistles emitted by a single individual, with no silence between the frequency contours (36)). We found that connected sequences contained SWs of different sub-categories indicating that dolphins are capable of emitting different variants of their own SWs (Fig.4a-d). Moreover, we observed connected sequences combining SWs of different individuals (Fig. 4e-f). These combined sequences could be used when an individual identifies itself and calls for another dolphin (e.g. I am Neo, and I am calling Nikita).

**Figure 4.**
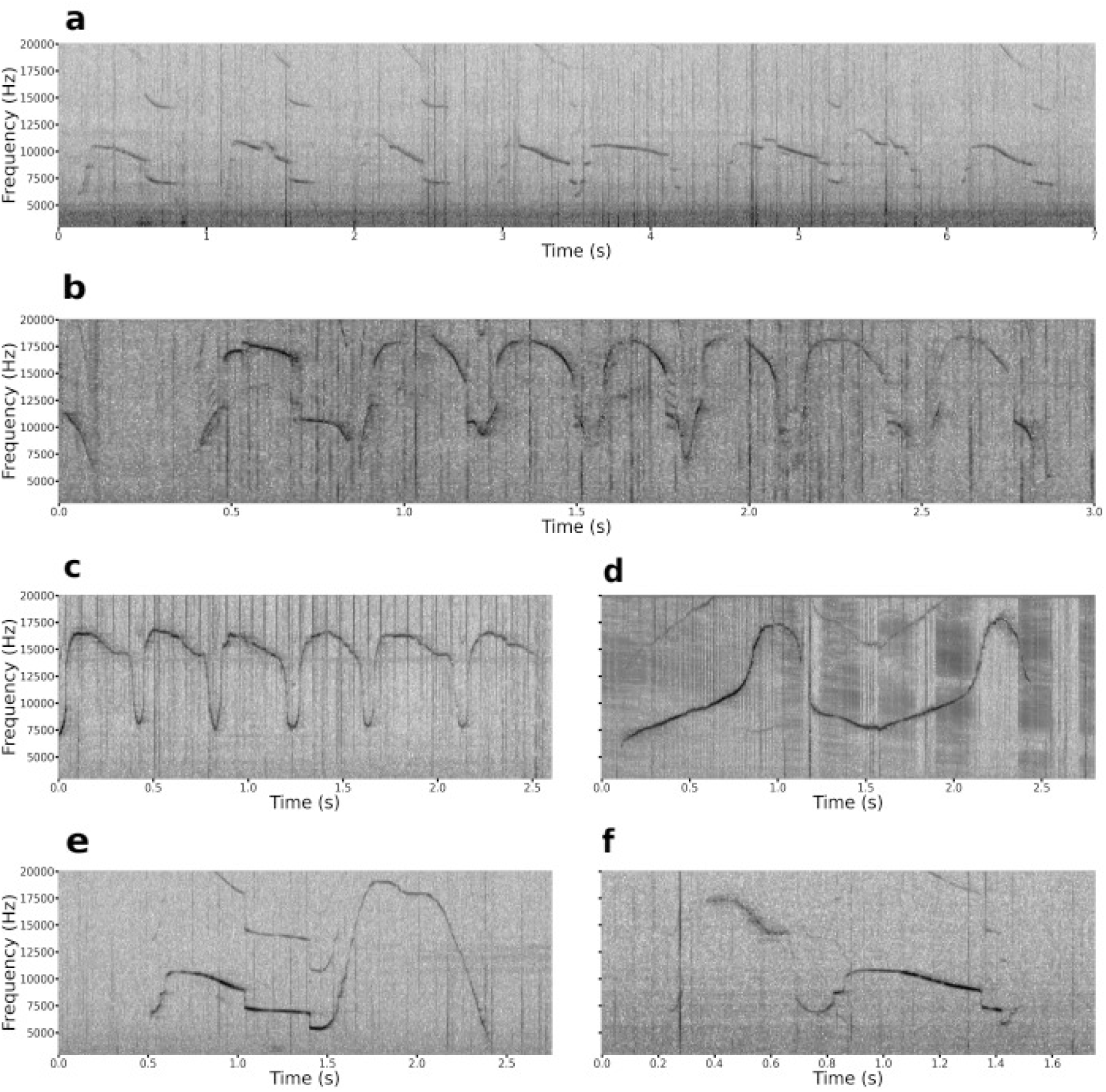
Connected sequences of whistles show that different SW variations are emitted by the same dolphin. a-d) Example spectrograms showing connected sequences of the SW_Neo (a), SW_Luna (b), SW_Nikita (c) and SW_Nana (d). A connected sequence indicates that all whistles were emitted by the same individual. e) As for (a), showing the connected sequence involving two SWs belonging to different individuals (SW_Neo followed by SW_Nikita). f) As for (e), containing the SW_Nikita followed by SW_Neo.

### The structure of social acoustic interactions

The acoustic interactions between dolphins were continuously recorded for one-hour periods, several times a day, over ∼5 months. Therefore, it was possible to study the interactions between the different whistle variants. The patterns that emerge from these interactions can provide insight into the social structure of the dolphin pod and principles governing their acoustic communication even in the absence of additional context (e.g. McCowan et al., 1999 (37), Ferrer-i-Cancho & McCowan, 2012 (38)).

To that end, we analyzed the structure of the pairwise correlations using a Markov chain model. A Markov chain displays the interactions between the different whistle variants (sub-categories), where the sub-categories are represented by nodes. These are interconnected by arrows. Their direction and thickness represent the probability of a given node to predict another (e.g. if sub-category A is emitted, what are the chances that sub-category B will be emitted within a given time window). To build the Markov chain model, we first computed the intervals that indicate whether two whistles belong to the same sequence, are considered as interactions, or are emitted independently from each other. For this purpose, we used a data-driven approach based on the distribution of inter-whistle time intervals across all the extracted whistles (Supplementary Fig.3a, see Methods).

To determine the statistical significance of these transitions, we used a statistical comparison with data-based null models (Supplementary Fig. 3b-f, Methods).The Markov chain model revealed that the structure of the network was organized according to several highly interconnected, large clusters. These clusters mainly included nodes belonging to the same main whistle category, an observation further supported by a community detection analysis (Louvain algorithm based on maximization of modularity, see Methods). Communities or “modules” are defined as groups of nodes strongly interconnected, and weakly connected with other groups. Our analysis identified a total of 5 communities. Four of these communities were composed of SW_Neo, SW_Luna, SW_Nikita and NSW_9 sub-categories. An additional one was composed of both SW_Nana and SW_Yosefa sub-categories (Fig. 6a). To quantify the division of the nodes into communities, we calculated the modularity of the network. Modularity is a measure of structure in a network ranging from -0.5 to 1: when modularity is positive, it indicates the presence of communities in the network (see Methods). The modularity value of the network of whistle sub-categories was 0.65, corroborating the existence of communities in the network of whistle sub-categories.

It is important to note that the nodes indicate the whistle sub-categories without information about the identity of the dolphin that emitted the whistles. However, SWs dominate the acoustic vocalizations of dolphins both in the wild and in captivity, representing between 50 and 90% of their whistle repertoire (25–27). In our entire dataset, the SWs represented 91.3% of the total number of extracted whistles. In our smaller emitter-labeled dataset, Neo produced its own whistle 53.5% of the time, Yosefa 83.4% and Nikita 95.5% (we did not have enough data to quantify the SWs of Luna and Nana). Therefore, we assumed that the main categories of SWs in the Markov model (communities) most probably represented sequences emitted by the owner of the whistle. However, a few individual nodes (whistle sub-categories) were found within communities belonging to a different SW (SW_Nikita (31) and SW_Nikita (36), Fig.6a). The latter may represent dolphins emitting SWs of other individuals.

Given that the history of the pod is well documented (34, 39) (Supplementary Fig.5a), we interpreted the results from the network visualization and analysis in light of their social structure. In our dataset, the SWs of Neo and Luna were the most frequent ones (Luna: 2980, Neo: 2260, Nikita: 1007, Yosefa: 627, Nana: 517). Indeed, Luna is the leader (dominant individual) of the pod, while Neo is the only male of the pod (Supplementary Fig.5a). Furthermore, the community analysis showed that nodes belonging to SW_Neo and SW_Luna did not appear in other communities (Fig.6a). This may indicate that Neo and Luna tended to produce monologue-like whistle sequences, while Nikita, Nana and Yosefa whistle in response to other dolphins, or no other dolphins produced their whistle (i.e. they are seldom called).

When we analyzed the compactness of the communities (the ratio between the average transition probabilities between nodes of a given cluster, and those with the rest of the nodes, see Methods), we found that the SW_Neo sub-categories were the most compact ones (SW_Neo = 24.35, SW_Luna = 16.23, SW_Nikita = 12.87, SW_Yosefa = 11.96, SW_Nana = 10.49). The SW_Yosefa sub-categories were one of the least compact communities, and appeared in the very middle of the graph (Fig.5a). The latter was quantified by measuring the distance between the center of mass of each SW main category to the center of mass of the whole graph. We found that SW_Yosefa sub-categories displayed the lowest distance (SW_Neo = 9.39; SW_Luna = 6.72; SW_Nikita = 6.04; SW_Nana = 5.08; SW_Yosefa = 3.56). This may indicate that Yosefa, rather than generating long sequences, emits her SW whenever other dolphins emit a whistle. Yosefa is a solitary *Tursiops aduncus* female dolphin who temporarily joined the pod of the Dolphin Reef for a few months during the recording period. One possible interpretation is that Yosefa made an effort to interact with the other dolphins to integrate within the pod. Alternatively, Yosefa could be using a different communication strategy to that used by the local dolphins. Indeed, personal communication from the divers at the Dolphin Reef confirmed that despite spending several months at the Dolphin Reef, Yosefa did never fully integrate into the pod.

**Figure 5.**
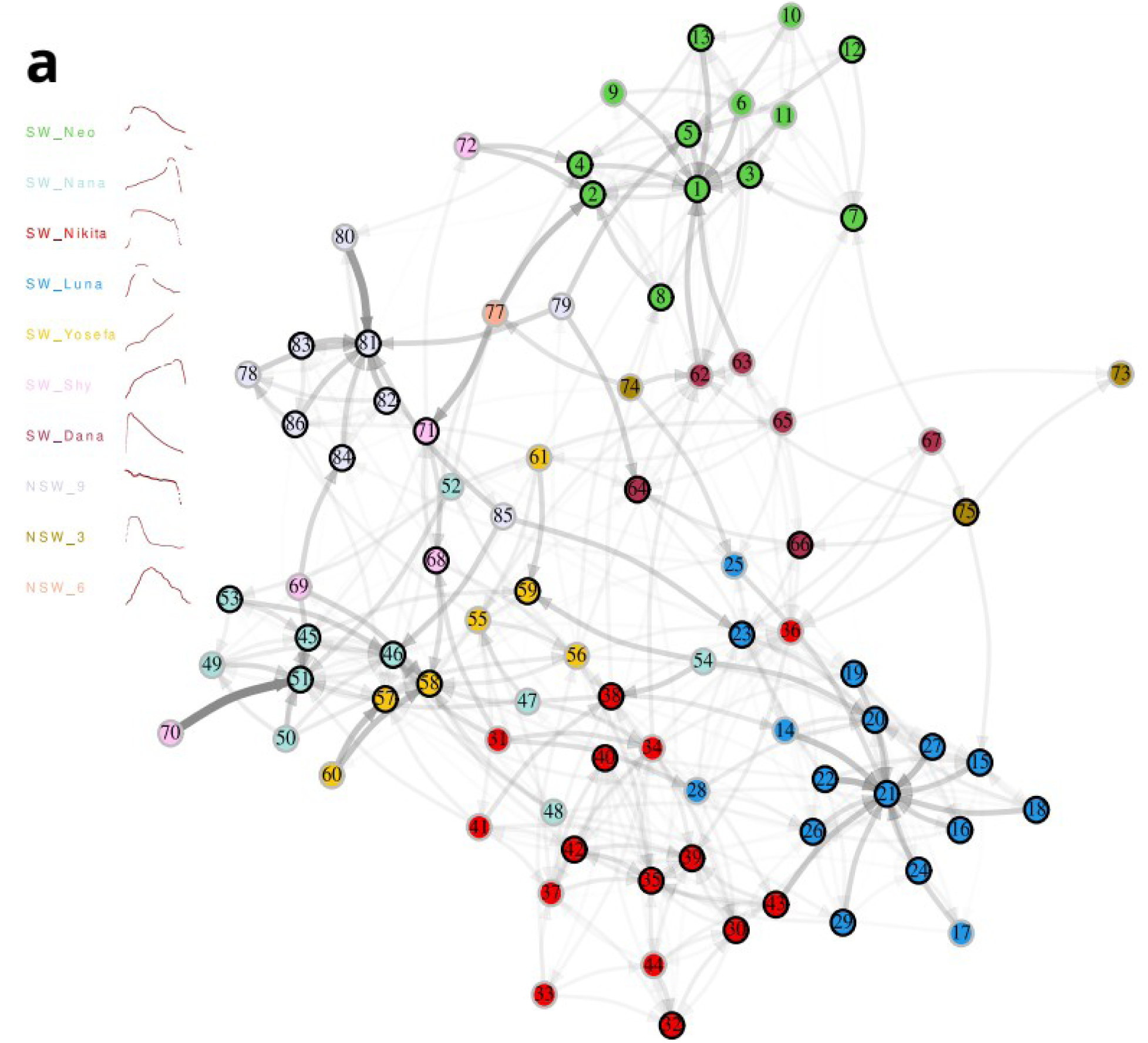
Markov Chain Model and hypergraphs capture the acoustic communication structure of the dolphins’ pod. Visualization of a Markov Chain Model where each node represents a whistle sub-category, and the arrows indicate the existence of a significant transition probability between two nodes. The thickness of the arrow is proportional to the transition probability. Nodes with black circles show auto-loops. Node colors represent the main whistle category (legends). Note the clustered distribution of the graph according to the whistle main categories (SWs and NSWs).

To corroborate this hypothesis and test whether the acoustic interactions between Yosefa and the rest of the pod evolved during the recording period (> 5 months), we built a Markov model for the first half of the recordings (10th of November 2019 to the 10th of January 2020), and a second one for the second half (11th of January to the 12th of March 2020, see Methods). We then analyzed the community compactness for the two datasets. We found that the compactness of the clusters of both graphs were not significantly different (p > 0.05, paired t-test). This result indicates that the acoustic interactions of Yosefa with the other individuals of the pod did not change during the period she spent in the reef. Furthermore, this suggests that the acoustic interactions between all the dolphins in the pod were stable across time.

A highly compact cluster of nodes from the category NSW_9 was found relatively isolated from the network (average distance with the rest of the clusters = 2.76±0.14; Compactness ratio = 37.19). This suggests that these particular NSWs are weakly related to other types of whistles. In contrast, we found that they are associated with burst-pulse sounds (Supplementary Fig.4).

When observing the Markov model (Fig.5a), we found that the nodes 62 to 72, belonging to two different NSWs (NSW_1 and NSW_2), were located in between the communities of nodes representing the SW of the different dolphins. Nodes 62 to 67 were located between the communities of Neo and Luna, while nodes 68 to 72 were positioned next to those of Neo and Nana. Given their peculiar position, we compared these nodes with those of deceased dolphins that once were part of the pod (Supplementary Fig.5a) (34, 39). For this, we computed the DTW distance between the unknown NSWs with the templates of the SWs of their ancestors (Supplementary Fig.5c). We found that these nodes closely matched the SWs of two female dolphins: Dana (died in 2013, mother of Neo and Luna) and Shy (died in 2015, mother of Nikita and Nana) (Supplementary Fig.5a). The SWs of other 17 dolphins that once were part of the pod (34), were never detected in our recordings. The SW_Dana sub-categories were found between the communities of Neo and Luna, while the SW_Shy sub-categories were placed close to Nana. SW_Shy sub-category 72, was also very close to the community of Neo. The latter may be the consequence of Neo being Shy’s sexual partner (Fig.5a). This is also supported by the community detection analysis (Fig.6a): nodes of SW_Shy were within the community mainly composed of SW_Neo and within the community mainly composed of both SW_Nana and SW_Yosefa. The nodes of SW_Dana were present in her offsprings’ community (Neo and Luna). Neo’s category was found in proximity to the nodes of both Dana and Shy. Divers in the dolphin reef indicated that Neo usually emitted the whistles of Dana and Shy prior to sexual interactions (personal communication). The finding that dolphins can emit the SWs of deceased individuals was also previously observed by A. Kershenbaum, T. Morgan and N. Shashar (personal communication).

**Figure 6.**
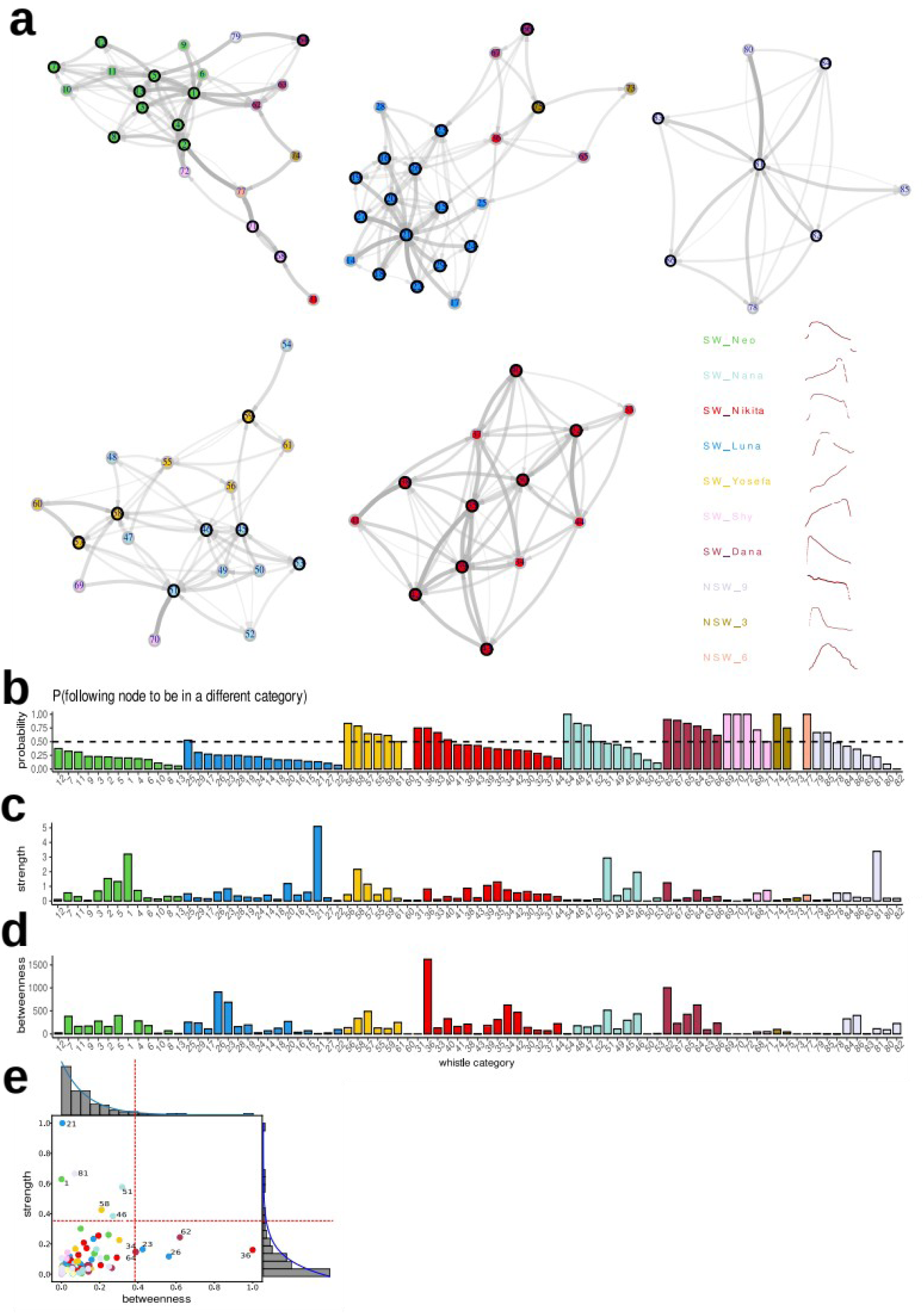
Network analysis of the dolphin acoustic interactions. a) The 5 resulting communities from the Markov chain model in Fig.5a. Each node represents a whistle sub-category, and the arrows indicate the existence of a significant transition probability between two nodes. The thickness of the arrow is proportional to the transition probability. Nodes with black circles show auto-loops. Node colors represent the main whistle category (legends). Note that most of the communities contain a single whistle sub-category. b) Barplot indicating for each whistle sub-category (node), the probability of outbound transition. Dashed horizontal line: probability = 0.5. The bars are colored according to the whistle category they belong to. The bars are all ordered by main whistle category, and in probability decreasing order within the main whistle categories. Note the absence of nodes with high probabilities for Neo (only male) and Luna (leader). The other SW categories only have a handful of nodes with high probabilities. c) As in (b), for the value of strength. The nodes are ordered according to (b). Note that only one node per main whistle category shows high strength values, mainly for Luna and Neo. d) As in (c), for the value of betweenness. Note that only one Nikita shows a single node with high betweenness value. e) Scatter-plot between strength and betweenness values. Every dot represents a single node. The color indicates the main whistle category of the node as in (a). The barplots represent the distribution of strength (y axis) and betweenness (x axis). The blue lines represent the fitted curves, determined by fitting exponential decay functions to the data. The red-dashed lines indicate the 90^th^ percentile of the data. Note that several nodes show high strength with low betweenness, and vice-versa.

### Network analysis of the acoustic interactions

Network theory provides a framework for understanding the structural and organizational principles underlying the connectivity of nodes within a network. The analysis of these connectivity patterns may shed light on the functional roles that different nodes play in facilitating the flow of information within the network.

Specifically, we computed for each node of the network (SW sub-category), three network measures: strength, betweenness and turn-taking probability. Strength reflects the level of connectivity of a given node with the rest of the network. It is defined as the sum of the weights of edges connected to one given node (we considered only inward connections). Nodes with high strength represent hubs that are vital for distributing information across the network (nodes that occur frequently, without necessarily connecting different communities). Betweenness captures the number of times a node is part of the shortest path between two other nodes (bridging between nodes), indicating the connection level of a given node to the network. They correspond to whistles that do not necessarily appear often, but they are fundamental to create connections between communities (nodes that assure communication between distinct, otherwise disconnected parts of the network) (40, 41). Turn-taking is the probability that for a given node, the following one will belong to a different whistle main category (Fig.6b). The latter will represent SW sub-categories emitted with the purpose to change the identity of the emitter dolphin. Using these 3 network properties, we found that all nodes of SW_Neo and SW_Luna showed turn-taking probabilities below 0.5, indicating that these vocalizations tend to be produced within a sequence of the same main category. In contrast, certain nodes from the other main categories displayed probabilities above 0.5. Among the nodes of SW_Nana, SW_Nikita and SW_Yosefa, several showed high turn-taking probabilities, indicating that these dolphins generate more individual whistles followed by the SW of another individual, rather than long sequences composed of the same SW main category. To quote an example, NSW_3 and NSW_6 have exclusively high turn-taking nodes. Thus, they could be specifically used to indicate the transition from one category to another (similar to the word ‘over’ in radio communications). Most of the nodes within the sub-categories of SW_Dana and SW_Shy showed high turn-taking probabilities. This may be the consequence of dolphins incorporating these SWs within their own sequences. In certain cases, the nodes with a high probability appeared closer to the nodes of a different category (node 36 and 31, Fig.6a). Thus, they could represent SWs emitted by other dolphins rather than the SW’s owner (node 36 could represent the SW_Nikita emitted by Luna, Fig.6a). For example, these vocalizations could have been produced by Neo and Luna in combination with their own SW (e.g. Nikita calling Neo or vice-versa, Fig.4e-f).

Strength analysis revealed that just one hub was observed by whistle category, and especially noticeable for SW_Luna (the leader, node 21) and SW_Neo (the only male, node 1). This suggests that only individuals high in the social structure (leader of the pod and only male) displayed hub nodes. The fact that there is only one within the individual’s repertoire may suggest that these nodes have a central functional role in the pod’s acoustic interactions. It is also possible that they use these hub nodes to send information without expecting a response (e.g. communicating orders).

The analysis of betweenness showed a different behavior. It was mainly relevant for SW_Nikita (36). This suggests that using this node, Nikita interacts with the entire pod. Despite Nikita’s lower hierarchical social role within the pod compared to Neo or Luna, this bridge node allows her to interact with all individuals of the pod. Nikita seems to be an individual with a cohesive function within the pod capable of distributing information across all individuals. Correlation analysis between strength and betweenness showed that most of the nodes displayed correlated levels of low strength and low betweenness. However, several nodes showed either high or low strength/betweenness ratios (high: 21, 1, 81, 51, 58, 46 and low: 36, 62, 26, 23, 34, 64). These nodes may play specific functional roles (e.g., transmitting orders vs. distributing information). The finding that nodes have different functional properties also supports the idea that SWs sub-categories represent potential linguistic units of dolphin acoustic communication.

### The order of sub-categories within whistle sequences

In linguistics, syntax represents the principles underlying the combination of basic units (words, morphemes) to build sentences (e.g. the grammatical rules dictating the order of words in a sentence). To study whether the sequences of the whistle sub-categories follow a particular order, **w**e first asked whether subcategories occur in a specific position in sequences. Specifically, we calculated the distribution of the positions of each SW sub-categories, within the emitted sequences. These distributions were then compared to those of a data-based null model, created by shuffling the whistles’ position. We found that no whistle sub-categories occurred systematically at specific positions in the sequence (p > 0.05, Kolmogorov-Smirnov test, see Methods). In addition, we also examined the relationship between whistle sub-categories and their position within the sequences by testing whether any significant interactions between position and sub-category exist. We did not find any significant difference between groups (p>0.05, Kruskal-Wallis test). Therefore, these results suggest that the organization of whistle sub-categories within sequences does not follow a specific order. Yet, it is possible that sequences display specific combinations of whistle sub-categories. To test this hypothesis, we calculated the high-order interactions between nodes within the main whistle categories (interactions within sequences). For this purpose, we built hypergraphs using whistle sub-categories (Supplementary Fig.6, see Methods). To compute the significant interactions between nodes (edges), we used a null model created by randomly shifting the order of whistles within sequences (see Methods). Across all main categories, the zero-order and the first-order edges had the strongest weights, and at least one strong pair of whistle sub-categories. For all SW sub-categories, the hypergraphs showed several zero, first and second order interactions between the different nodes (Supplementary Fig.6). Although no clear order emerged in the sequence organization, the hypergraph analysis showed specific pairs and triplets that are more likely to occur together. The use of specific combinations of SW sub-categories suggests that dolphin acoustic communication follows certain rules.

## Discussion

Despite more than 70 years of dolphin research (42), our understanding of their acoustic communication remains limited. A significant obstacle has been posed by the difficulty to monitor, in natural conditions, the acoustic interactions from the same group of dolphins over long periods of time. Here, we studied dolphin acoustic communication in the Dolphin Reef, Eilat, Israel, where a previously studied pod of dolphins spends most of their time close to the shore (34), allowing for long-term recordings of their acoustic interactions, in a natural environment.

Using this unique large dataset, we investigated the variability observed in the frequency contours of whistles.Here we analyzed 8566 whistles from a known pod of dolphins living in a natural environment. We observed that the emitted whistles displayed distinct frequency modulations. These whistle variations could not be explained by a random process, nor the different identities of the dolphins emitting the whistles. In the case of the SWs, these variations involved stereotypical transient frequency modulations at specific time points of the frequency contours, and changes in the duration of the whistle. In most cases, although not exclusively, the modulations were observed at the maximal frequency (peak) and at the end of the frequency contours. In line with previous research that suggested the modulation of the SWs is context-dependent (31), we propose that this variability represents a strategy to encode additional information that goes beyond the dolphin’s identity. Specifically, we suggest that the frequency envelope of the SW may function as a carrier signal (e.g. the broadcasting frequency of a radio station), within which the transmitted information is encoded using specific frequency or amplitude modulations. This strategy would enable the receiver individual to know the identity of the dolphin transmitting the information, even if it is out of sight. These carrier signals could also function as acoustic communication channels (each SW would be a different channel), enabling simultaneous “conversations” between dolphins in a pod, without interfering with one another. This hypothesis conciliates the contrast between the dolphins’ complex social structure, which requires an advanced communication system, and their limited whistle repertoire, mainly dominated by SWs. This idea is also supported by a previous study showing that whistle variations could be associated with emotional states (31). It is important to mention that beyond whistles, dolphins also emit burst-pulse sounds believed to transmit emotional states (43). Experiments involving underwater cameras to assess the context of the emitted whistles (e.g. agonistic, courtship, fishing, or play behaviors) would provide further ground to support these hypotheses.

The classification of whistles in sub-categories (units), in combination with a Markov chain model enabled us to study the structure of the acoustic interactions between dolphins in the pod.

Network analysis of the pod’s acoustic interactions showed that network communities are mainly composed of whistle sub-categories belonging to the same main whistle categories (same SW). Under the assumption that the emitters mainly generate their own SWs, this result suggests that individuals produce sequences using different sub-categories of their SWs (nodes). Within these sequences, we found that certain nodes have distinctive functional roles (such as turn taking, hubs and bridges). Turn taking nodes predict the change of main whistle category within a sequence. Hubs characterize a tendency to monologues, while the presence of bridges identifies nodes that are involved in interactions across the entire network. We observed that not all dolphins exhibit identical distributions of hubs and bridges. Interestingly, this distribution showed correspondences to the dolphins’ social role in the pod. For example, Neo (the only male) and Luna (the dominant female of the pack) were the only individuals showing hubs, while Nikita, was the only one showing bridge-like nodes, highlighting her role as an individual who is crucial to the pod cohesion.

These results allowed us to draw conclusions about the pod’s social organization leveraging only their acoustic productions. Moreover, the non-random structure of the acoustic interactions, and the fact that different SW sub-categories have different functional roles (they are not interchangeable), validate the sub-categories as potential linguistic units and further support the hypothesis that the observed variability of the whistles carries information beyond the dolphins’ identity.

The analysis of the structure of the SW sequences did not detect fixed-order associations that go beyond random juxtaposition of single nodes. However, using hypergraphs we found statistically significant frequent pair and triplet associations between nodes. This suggests that a combinatorial structure of nodes could exist, as the lack of fixed-order in sequences does not rule out the existence of a communication system governed by rules. Indeed, in chimpanzees, the order of combinations of vocalizations is not always strict, but combinations exist nonetheless as they are produced non-randomly, and in distinctive contexts (44–46). In certain human languages (Warlpiri, O’odham, Mohawk) the order of words in a sentence is grammatically irrelevant but grammar is still present (free word-order grammars) (47–49).

Previous studies suggest that some animals display the ability to remember past or deceased individuals. For example, elephants show interest in visual encounters with the skills of deceased kins (50), and ravens seals recall the vocal calls of previous group members or kins, even after several years of separation (51). Nonetheless, they do not show the ability to transmit the identity of the deceased individuals employing an acoustic signal. Here, we found that the dolphins emit the SWs of their deceased mothers. A previous study showed that dolphins have the cognitive capacity for referential use when trained to do so, by pressing a ‘yes or no’ paddle to answer whether an object was present or not in their tanks (52). In our study, we observed that dolphins emit the identity of individuals that are no longer present in place and time, demonstrating that dolphins indeed use displacement and referentiality. The combination of displacement, referentiality and arbitrariness (no relationship between the sound and meaning when emitting SWs (53)) has currently been found only in human language. This result also raises the question whether previously reported NSWs, that represent just a small portion of their acoustic repertoire, could just encode the SWs of deceased mothers rather than carrying contextual or emotional state information, as previously hypothesized (31). In this case, the vocalization repertoire would be limited to burst-pulse sounds and whistles. The latter would only represent ‘names’ of living dolphins or deceased mothers, while additional information may be transmitted by their modulation. Overall, our results provide new insight and principles underlying the complexity of dolphin acoustic communication necessary to maintain a fission-fusion society. The future addition of well-detailed behavioral context and emitter identity to the acoustic datasets will be an important step towards ultimately deciphering dolphin acoustic communication.

## Methods

### Dolphins’ environment

The datasets were acquired at the Dolphin Reef, in Eilat (Israel), a touristic beach located in the northern part of the gulf of Aqaba. The Dolphin Reef is the natural habitat of a pod of bottlenose dolphins that spend most of their time in the proximity of the shore. The dolphins can leave and enter the reef without any restrictions (Supplementary Fig.5b). At the reef, dolphins may spontaneously interact with humans. These interactions are only initiated by the dolphins. The dolphins are not trained or incentivized to engage in any type of interaction or behavior. The pod of dolphins, at the time of the recordings was composed of five individuals: Nikita (female, 17 years old), Luna (female, 20 years old), Nana (female, 25 years old), Yosefa (female, age unknown) and Neo (male, 15 years old). All individuals were born within the Dolphin Reef, except for Yosefa. More information about the family tree of the four dolphins (*Tursiops truncatus ponticus*) can be found in Perelberg et al., 2010 (34) (Supplementary Fig.5a). The signature whistles of these four dolphins were determined in previous studies (39, 54). Yosefa (*Tursiops aduncus*), is a lonely female from outside the dolphin reef, from the Indian Ocean. Yosefa, was observed for the first time at the Dolphin Reef on the 28 of November 2019, and sporadically joined the pod during the recording period. It has been shown that the *Tursiops aduncus* also produces signature whistles (12). Yosefa’s signature whistle has been identified in the present study using the SIGID method (Signature Identification) (55).

Each individual presents physical characteristics that allow their identification. Specifically, these idiosyncrasies are observable at the level of the dorsal fin and the jaw, which show different spots and shapes for every individual, as well as the shape of the melon, tone of the skin and size.

### Acquisition, detection and extraction of whistles

Acoustic data was acquired using a 8104 Brüel & Kjær® hydrophone, installed at the entrance of the reef (Supplementary Fig.5b). The hydrophone was connected to a 1704 Brüel & Kjær® pre-amplifier (100x gain) and a Spectra DAQ-200 acquisition device (acquisition frequency 1Hz-96kHz). The data was recorded on a Linux computer that controlled the acquisition times using *crontab* command. The acoustic signals were recorded everyday at fixed times: between 5:45 and 11:45, data was recorded every hour. Between 13:45 and 19:45, data was recorded every two hours. The recorded datasets were saved as *wav* files and transferred every night to a server at the Ecole Normale Supérieure. The datasets were recorded from November 1^st^, 2019 and March 12^th^, 2020. Extraction of whistles from the recordings was performed in two steps: i) an initial, manual annotation of a limited number of whistle contours (n=852). ii) a custom-built, automatic extraction algorithm based on spectral energy detection and dynamic time warping (DTW), using as templates, the annotated whistles.

For the manual extraction, the fundamental frequencies of 852 whistles were annotated using the Silbido tool on Matlab (56): this allowed us to identify the frequency and timestamp of each whistle. In order to optimize the subsequent automatic classification, all whistles were annotated individually, even those being part of so-called "multi-loops": whistle instances constituted of repetitions of the same elements at less than 0.5s intervals (12).

These whistles were then classified automatically, in order to identify a first inventory of whistle categories to provide the templates for subsequent automatic extraction and sequence analysis steps. For the classification, we used ARTwarp (35). ARTwarp uses an unsupervised learning algorithm, an adaptive resonance neural network, and DTW as a distance metric between the frequency profiles to cluster whistle contours with similar shape. The use of DTW ensures that frequency contours of similar shapes are clustered together despite their potential differences in the scale of their time series (e.g. similar whistles can be compressed or stretched).

The ARTwarp clustering resulted in the identification of 10 main categories of whistles: 5 signature whistles of the dolphins constituting the pod (SW_Neo, SW_Luna, SW_Nana, SW_Yosefa, SW_Nikita). Additionally, 5 whistles have been identified as non-signature whistles (NSW_1, NSW_2,NSW_3, NSW_6, NSW_9).

The extraction program (Supplementary Fig.1), developed in Python, is based on templates representing each of these clusters. The algorithm carries out the following steps:

i. Detection of frequency contours based on the variance of the frequencies at each time point.
ii. Reconstruction of the detected frequency contours using the frequency of maximal power intensity, for each time point.
iii. Calculation of the DTW distances between the frequency profile of the templates and those reconstructed from the detected whistle contours. The DTW distance is defined as the sum of the differences between every point associated in the DTW algorithm. We divided this value by the number of points of the reconstructed frequency contour.
iv. Extraction of the detected whistles if the DTW distances are below the threshold. The thresholds were defined manually and adjusted after observation of the preliminary results, to minimize false positives. This threshold ranged between 1150 Hz and 7000 Hz.
v. Identification of the beginning and end of the extracted whistles. Using the DTW points association between the template and the reconstructed whistle contour, the boundaries of the whistle were curated. Points of the observed whistle contour were dropped if a substantial number of points were linked to a single point on the template in the first and last 10% points of the model whistle.

Using this approach, we extracted 8566 whistles. The whistles from each main category were clustered a second time using ARTwarp, in order to detect possible conserved frequency modulations (signature whistle sub-types). The vigilance threshold value was adapted for each sub-category according to the method described by Deecke and Janik (35) for vocalizations whose biological categories are not known in advance. This method minimizes the variation within each category and maximizes the variation between categories. The vigilance values we obtained were: 85.0 for SW_Neo, SW_Nana, SW_ Yosefa, NSW_1, NSW_2, NSW_3; 84.44 for SW_Luna; 83.33 for SW_Nikita; 90 for NSW_9. Sub-categories containing less than 10 whistles were discarded.

This dataset is available at https://hub.bio.ens.psl.eu/index.php/s/zMZdy3ZgGW8CY38

The Python code used for the extraction of the whistles can be downloaded at https://github.com/zebrain-lab/Dolphins

Additionally, we created a sub-dataset of 402 whistles labeled with the identity of the emitter. This dataset was created by manually annotating video footage in which individuals were captured while swimming alone. This ensured we could label the sounds recorded at this time with the identity of the emitter.

### Probabilistic patterns in whistle sequences

To determine the maximal interval between whistles to be considered as sequence, (i.e. the limit between acoustic interactions and isolated vocalizations), we computed the inter-whistle time distribution of 4000 data-based null models (Supplementary Fig.3a, red). The null model distribution was generated from our data using the shifting method. This method involves randomly altering the time stamps of whistles in each recording. The shifting method eliminates any potential interdependence among the whistles while maintaining the overall count of vocalizations. The time values for shifting were randomly selected within the range of 0 to 1000 seconds.

We fitted both distributions with a Gaussian-exponential decay mixture function : 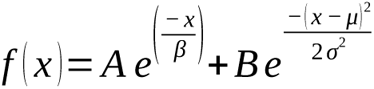 where A and B are the normalization constants, β the parameter for the exponential decay and (μ,σ) are the Gaussian parameters. We then identified the interval at which the null-model distributions intersected the dataset distribution (5.94 secs, Supplementary Fig.3a, black dashed line) to determine what constitutes a sequence: each whistle separated by an interval smaller than 5.94 seconds was considered as part of a sequence.

In a second step, we modeled the sequences obtained with this approach using a Markov Chain model. We started with the simplest and most ecological model, namely a first-order Markov Chain. Such models are widely employed in the early stages of analysis of a communication system (57, 58) as they are able to capture simple relationships between bigrams as well as higher-order relationships.

We built our model using sequences of 2 elements (whistles), and defined (i) a finite number of possible states, the sub-categories considered in the analysis (ii) the transition probabilities between these categories, gathered in a transition matrix. For each pair (i,j) of whistle sub-categories, the transition probability from i to j, denoted P_ij_, is the conditional probability that *i* is transmitted knowing that *j* has just been transmitted. These probabilities are estimated by the Maximum Likelihood, based on the number of occurrences of each transition (57).

To test the significance of each transition *i→j* and thus reveal pairs of whistles (*i,j*) whose co-occurrence in 2-element sequences can not be explained by a random process, we compared the transition matrix obtained from data to a set of 150000 data-based null models generated via the shuffling method. This approach consists in randomly changing the whistle sub-category in each recording so that the number of vocalizations is kept, while any potential dependence between them is lost. Direct two-tailed randomization tests were then performed for each of the possible *i→j* transitions. The p-value for each transition was estimated by directly comparing the observed P_ij_ to the distribution of P_ij_ in the null models (59). The observed *i→j* transition was interpreted as statistically significant if the p-value was lower than 5%.

A Markov Chain can be visualized in the form of a transition diagram, a weighted directed network in which: (i) the nodes correspond to the states considered in the model (ii) the directed edges represent the transitions between these states, weighted by their probabilities. Figures were produced using the *Igraph* package of *Rstudio*.

To characterize the structure of the whistles’ network, we computed node-level measures and global measures using *Igraph* and *tnet* packages from *Rstudio*. The analysis was based on the directed and weighted graph without disconnected elements, with the aim to keep information on the order of the tokens in the sequences.

Specifically, we calculated: strength centrality, i.e. the sum of edge weights connected to one given node. Betweenness centrality, defined as the number of geodesic paths (shortest paths) that pass through a given node. A “path” is typically a shorthand for “geodesic path” or “shortest path”— the fewest number of edges required to get from one node to another (60). These network measures may reveal hubs in the network (highly connected nodes or nodes involved in heavy-weight connections). The centrality of a node could indicate the node’s influence in the network. The more important a node will be in the communication between the nodes of the network, the higher its centrality measure will be.

In order to describe the structure of the social acoustic interaction, we studied “turn taking” dynamics. We computed, for every node, the probability of the following node to belong to a different main category after a transition. In other words, we considered the probability that after the occurrence of a certain whistle type the next whistles belong to a different main category.

To study the global structure of the network, we computed a community detection algorithm. We used the Louvain algorithm, based on maximization of modularity. Modularity evaluates the quality of a given partitioning. A high modularity value means that the network is structured in communities.

### Hypergraphs

To study the higher-order structure of the interactions between categories of whistles, we used hypergraphs. The hypergraphs capture how often a particular subset of whistles co-occur. We used the same threshold obtained from our inter-whistles time interval analysis to define a whistle interaction. The weights of the hyper-edges were scaled in proportion to the number of interactions between the individuals without taking into account the directionality of the interactions.

### DTW variability in frequency modulations

To perform a more detailed analysis of small modulations in the signature whistles, we developed a method to quantify these variations. The goal was to understand whether variability occurs uniformly across a whistle frequency contour, or on specific portions of it. The technique involves the use of DTW to align each instance of a signature whistle to a model whistle (a whistle not showing modulation) by category. Specifically, for each whistle the frequency contour was first smoothed using a Hanning window, and subsequently dynamically aligned to a model using DTW. This step ensured that whistles were normalized in the time-scale to allow focusing exclusively on frequency modulation patterns. Once a model whistle was selected, we performed time-series K-means clustering to identify modulation patterns. The number of clusters at this step was determined empirically and adapted to each whistle category. We then measured the Euclidean distance of each whistle to the model whistle, and calculated the average by cluster.

## Acknowledgments

We extend our sincere gratitude to the dedicated individuals who contributed to the success of this research endeavor. Special thanks to Anita Paparelli, Alexis Emanuelli, Advat Gal, Alex Sheremet, Hadas Zion, Taly Ouzen, Loyal Rauch, Eden Detooker, Sophie Donio, Frank Veit, Bryan Hansen, Guy Shoam, Roni Zilber, Natalie Chernihovsky and Nadav Solomon, AIH for their invaluable support. We thank Emmanuel Chemla, Joël Attia, Olivier Adam and Philippe Schlenker for comments on the manuscript.

This project was supported by the FIRE PhD program, funded by the Bettencourt Schueller foundation, and the EURIP graduate program (ANR-17-EURE-0012) to F. Mustun, the European Union’s Horizon 2020 research and innovation program under the Marie Skłodowska-Curie grant agreement No 945304 – Cofund AI4theSciences to C. Semenzin, and The CNRS International Emerging Actions (IEA, IEA00637) to G. Sumbre, N. Shashar and G. de Polavieja. Gonzalo G. de Polavieja and Dean Rance were also supported by the Champalimaud Foundation.

---

**Supp. Figure 1.**
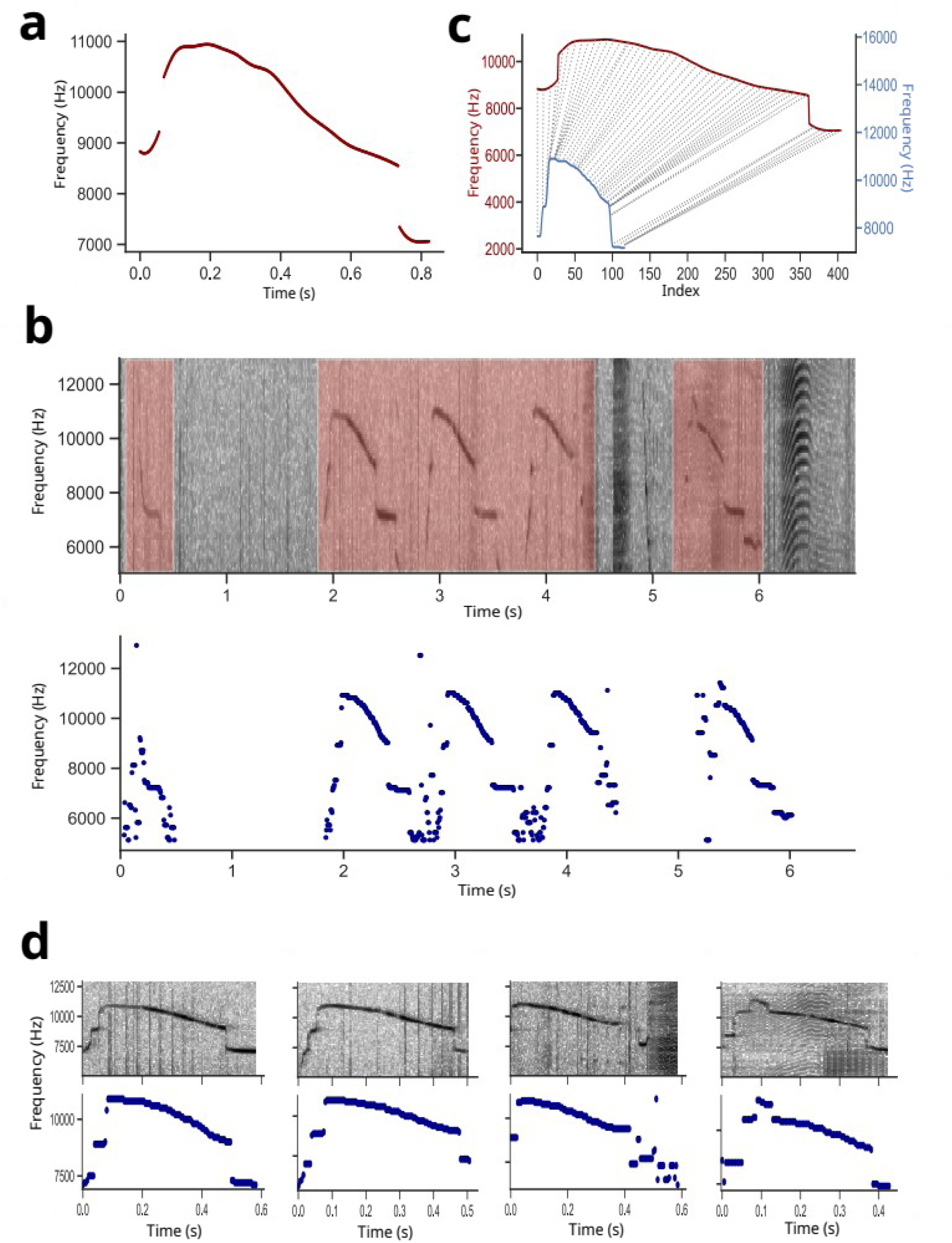
Method for the automated extraction of whistles. a) Frequency contour of the template whistle used for this example (SW_Neo). b) Top: example of spectrogram showing 5 whistles and 2 episodes of burst pulses. Red patches: automatically detected whistles, based on the frequency variance for every time bin of the spectrogram. Bottom: the frequency contours reconstructed from the spectrogram on top, by selecting the frequency with the maximum amplitude at each time point. c) Example of point association between the template whistle (red), and a detected whistle contour (blue) using the dynamic time warping algorithm. The dashed lines connect points reflecting the shortest distance between the two frequency contours. d) Four examples of extracted whistles. Top: spectrogram. Bottom: extracted frequency contours.

**Supp. Figure 2.**
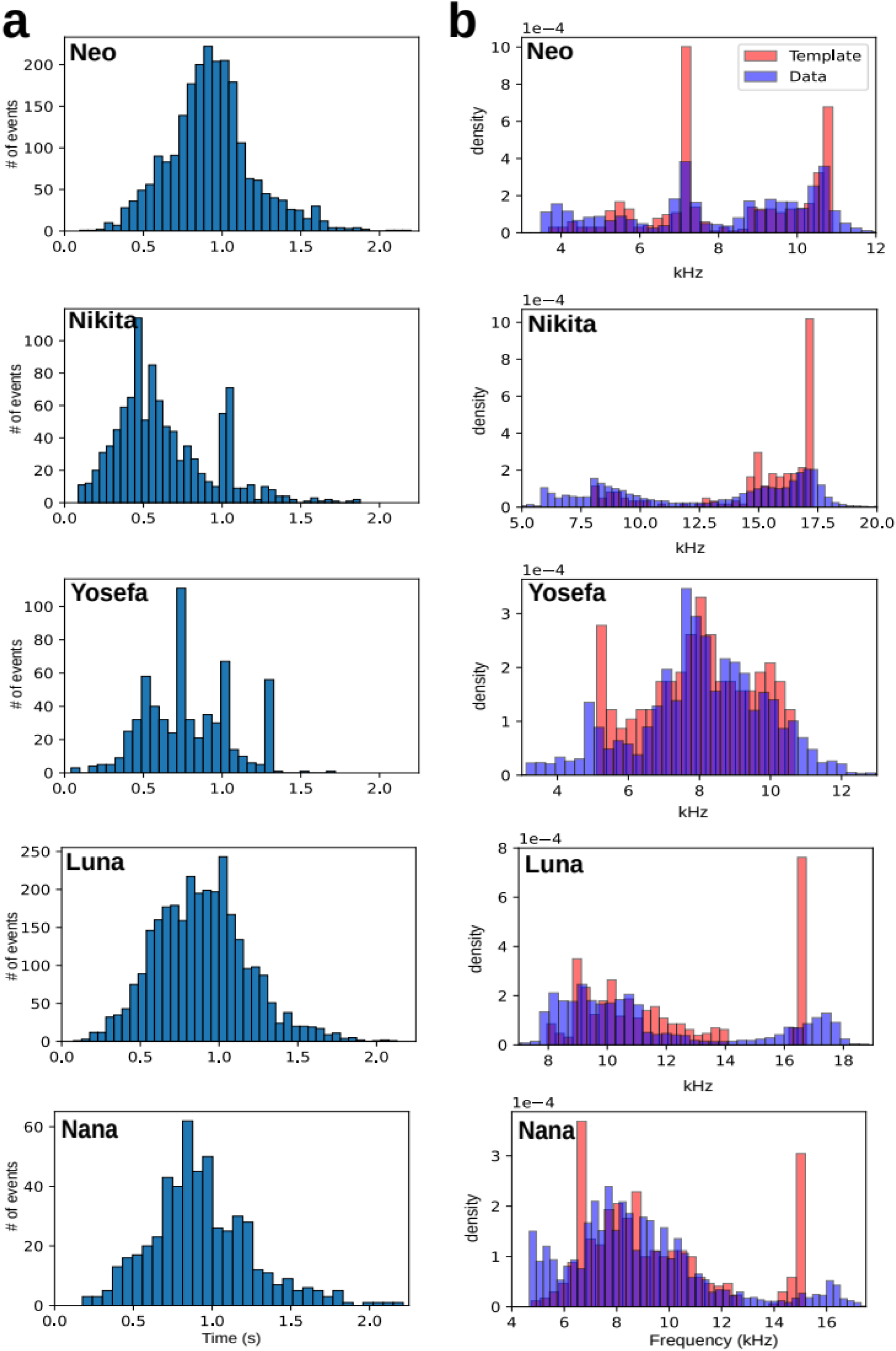
Distribution of duration and frequency of the SW main categories exhibit variability. a) Histograms depicting the SWs’ duration of each of the 5 recorded dolphins. Note the large distribution across all the SWs, indicating that dolphins modulate the duration of their SWs. b) Normalized distribution of the frequencies composing a template of the SWs of each dolphin (red). Normalized distribution of the frequencies composing the SWs in the data (blue). Note that the data distributions appear spread out around the peaks of the templates, indicating that dolphins use frequency modulation when emitting the SWs.

**Supp Figure 3.**
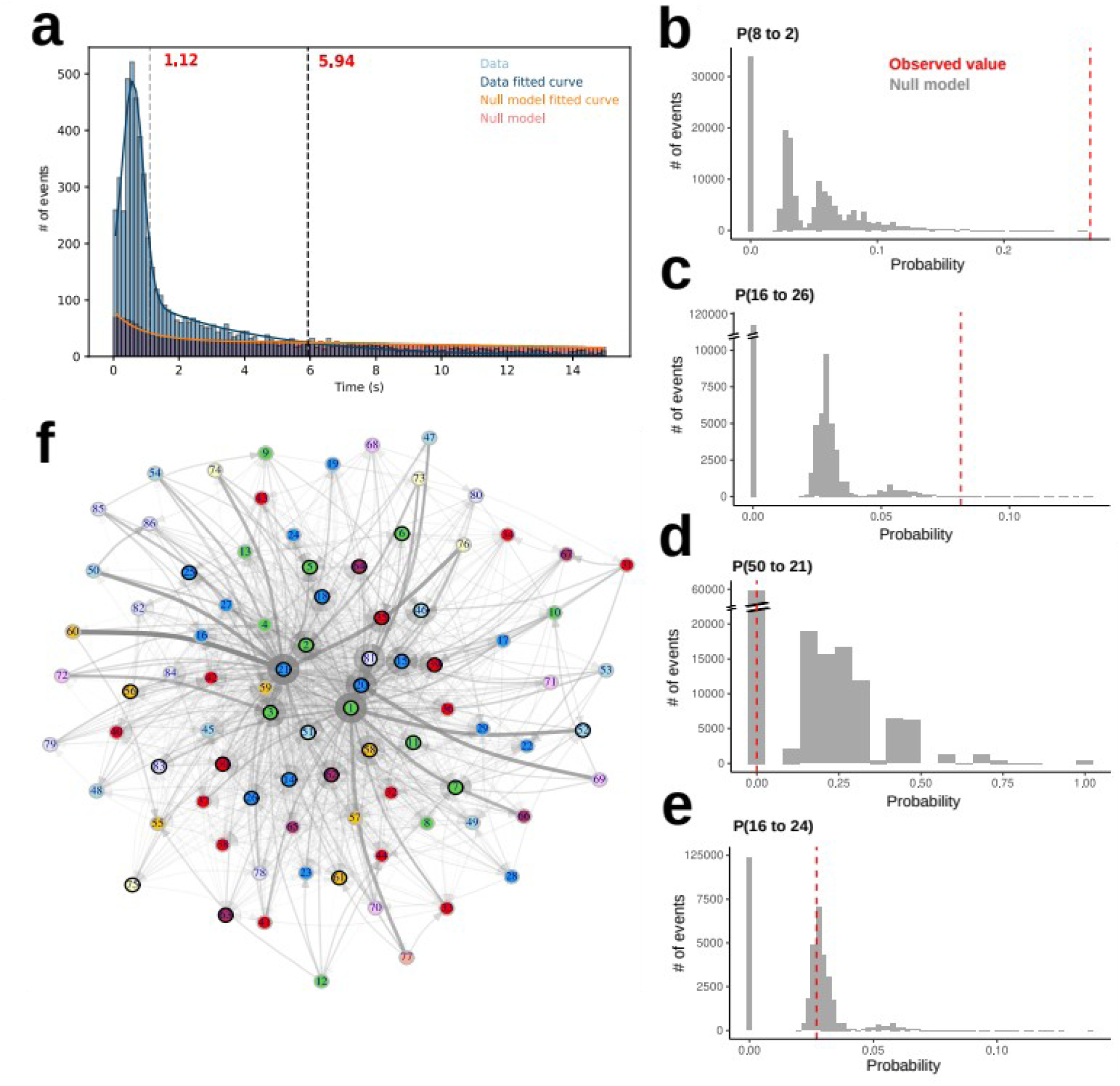
Statistical analysis of the interactions between the whistle sub-categories. a) Distribution of inter-whistle time intervals from the data (blue), and from the random null model (red). A Gaussian and exponential decay mixture function was fitted to the two distributions. Black vertical-dashed line (5.94 s): intersection between the data distribution and that of the null model, indicating the maximal interval between acoustic interactions. b, c) Examples of two probability distributions between two nodes (top: 8 to 2, bottom: 16 to 26) observed in the null models. Red-dashed vertical line: the observed probability in the dataset. The latter was statistically compared to the distribution of the null model. The probabilities in the dataset were significantly different from those in the null models (p<0.001, and p=0.004, direct randomization test). d, e) As for (b,c) for transitions probabilities between nodes 50 to 21 and 16 to 24 which were non-significant (p=0.543, and p=0.233). f) Example of a random network obtained by shuffling the order of whistles within sequences. Note the lack of clustered nodes of the same main category (same color) as seen in the observed dataset network.

**Supp. Fig 4.**
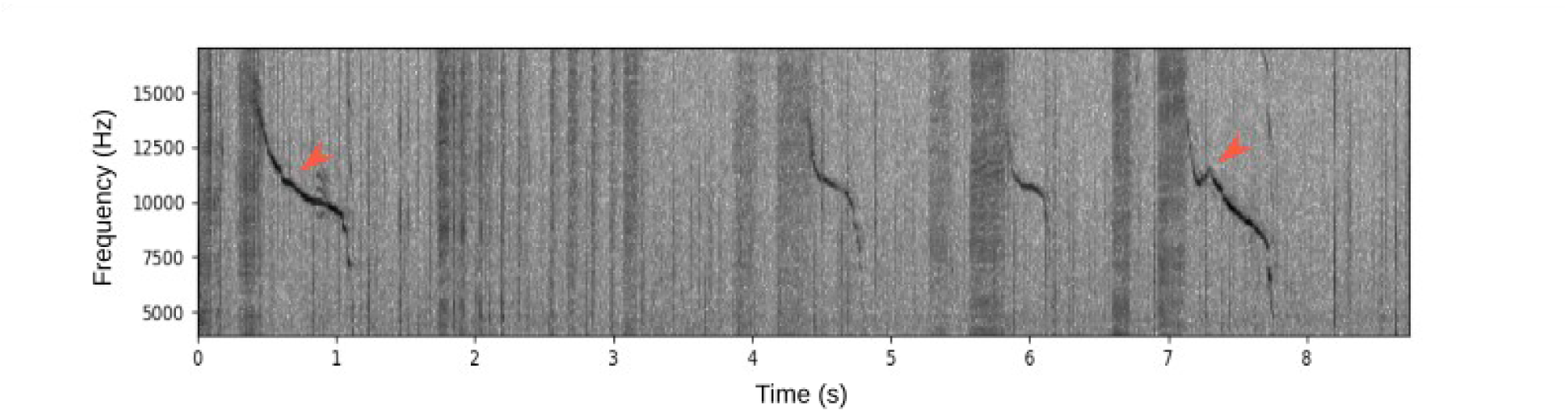
Non-signature whistle category 9 (NSW_9) is associated with burst-pulse sounds. A spectrogram showing an example of a sequence of four NSW_9 belonging to different sub-categories. NSW_9 is consistently preceded by a burst-pulse sound independently of the sub-categorization. Note the variations in duration and frequency contour (red arrows) of the whistle.

**Supp. Fig 5.**
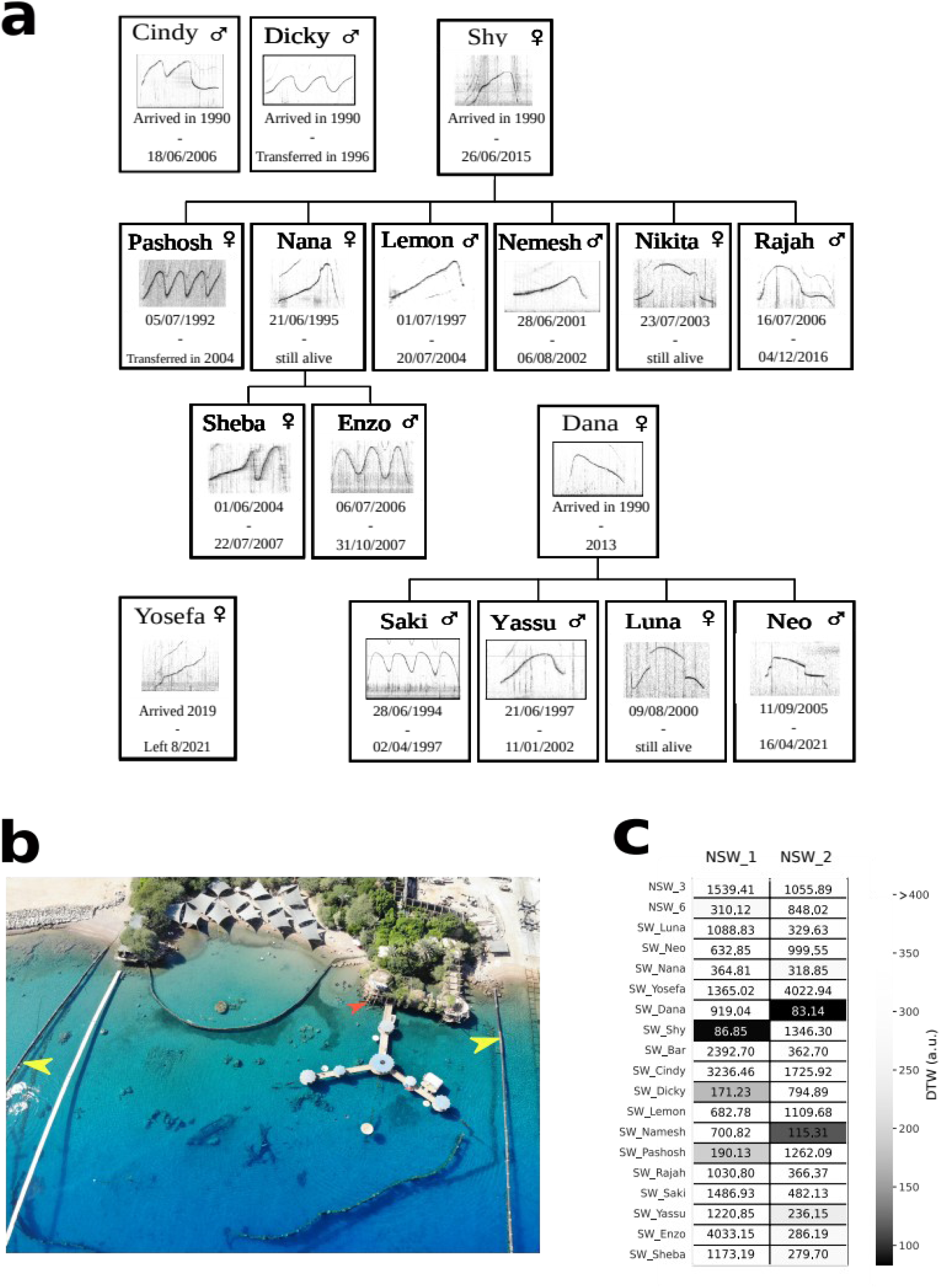
Dolphins emit the SWs of their deceased mothers. a) Family tree of the dolphins belonging to the pod, showing their sex and SWs. Cindy, Dicky, Shy and Dana arrived at the Reef in 1990 and Yosefa is a wild dolphin that joined the pod during the period of our recordings (from 01/10/2019 to 12/03/2020). Dicky and Pashosh were transferred back and released in the Black sea in 1996 and 2004 respectively. b) An aerial photo of the Dolphin Reef (Eilat, Israel) taken by David Tom Torjman. The red arrow shows the placement of the hydrophone used for the recordings. The yellow arrows indicate the surface barriers that prevent boats from entering the area, but dolphins can pass through to access the open sea. c) Matrix showing the DTW distances between the NSW_1 and NSW_2 with the signature whistles of all the dolphins that are or were part of the pod. Note that SW_Dana and SW_Shy pop out from all the rest.

**Supp. Fig 6.**
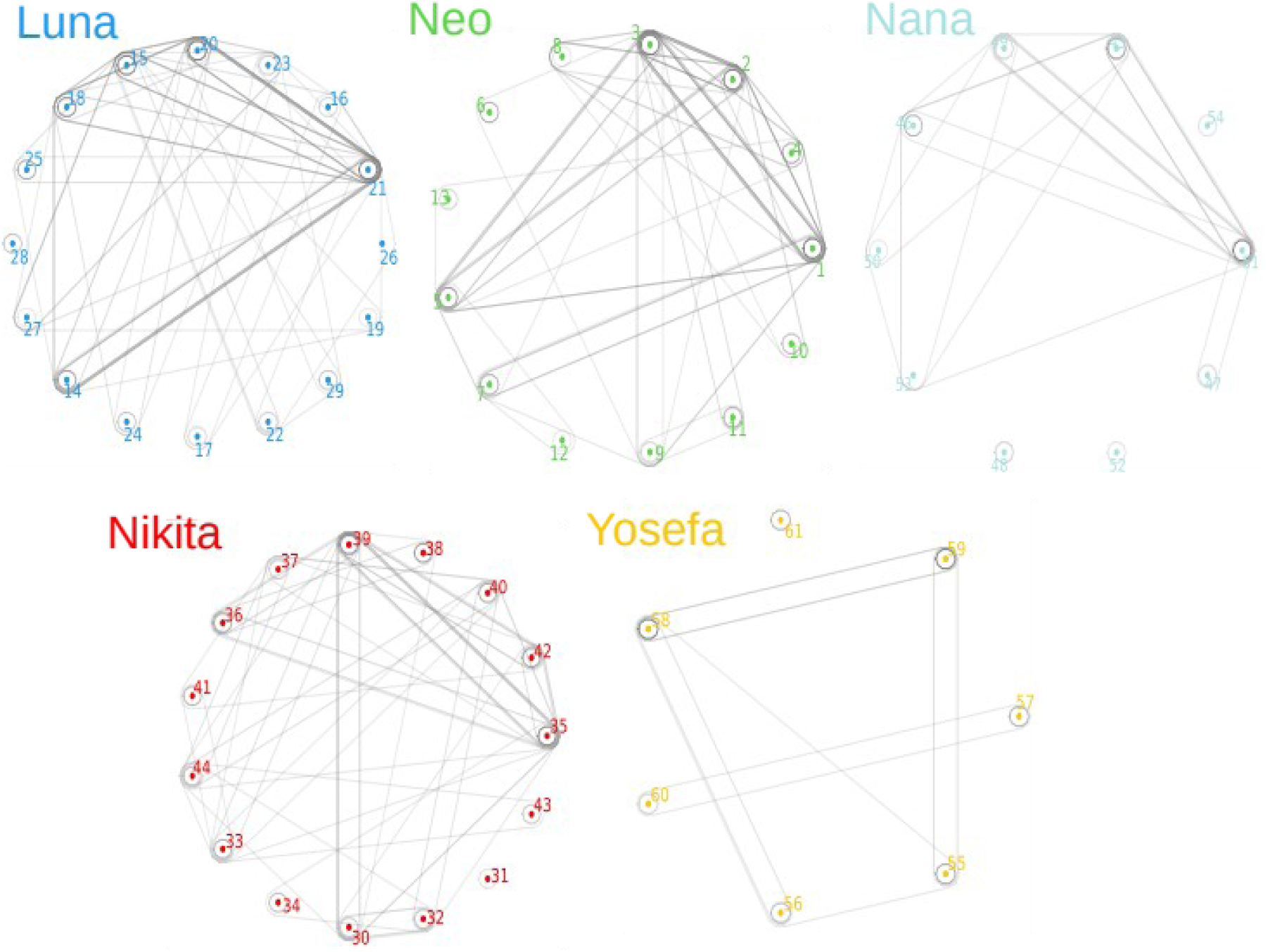
Hypergraph analysis of the intra-communities structure shows combinations of SW subcategories. 5 hypergraphs built only using the edges within the same main whistle category (one hypergraph for each dolphin). All zero-, first-and second-order co-occurrences were taken into account. Note the co-occurrences of specific SW sub-categories (nodes) within communities, suggesting that during sequences, dolphins emit specific combinations of SW sub-categories.

